# Biomolecular Condensation Drives Leukemia Caused by NUP98-Fusion Proteins

**DOI:** 10.1101/2020.11.16.384271

**Authors:** Stefan Terlecki-Zaniewicz, Thomas Eder, Johannes Schmöllerl, Theresa Humer, Natalie Kuchynka, Katja Parapatics, Elizabeth Heyes, Fabio G. Liberante, André C. Müller, Florian Grebien

## Abstract

NUP98-fusion proteins cause acute myeloid leukemia via unknown molecular mechanisms. All NUP98-fusion proteins share an intrinsically disordered region (IDR) featuring >35 repeats of Phenylalanine-Glycine (FG) in the NUP98 N-terminus. Conversely, different C-terminal NUP98-fusion partners are often transcriptional and epigenetic regulators. Given these structural features we hypothesized that mechanisms of oncogenic transformation by NUP98-fusion proteins are hard-wired in their protein interactomes. Affinity purification coupled to mass spectrometry of five distinct NUP98-fusion proteins revealed a conserved set of interactors that was highly enriched for proteins involved in biomolecular condensation. We developed biotinylated isoxazole-mediated condensome mass spectrometry (biCon-MS) to show that NUP98-fusion proteins alter the global composition of biomolecular condensates. In addition, an artificial FG-repeat containing fusion protein was able to phenocopy the induction of leukemic gene expression as mediated by NUP98-KDM5A. Thus, we propose that IDR-containing fusion proteins have evolved to uniquely combine biomolecular condensation with gene control to induce cancer.

AML, NUP98, fusion protein, AP-MS, LLPS, biCon-MS, condensate

## Introduction

Cancer-associated chromosomal rearrangements often result in the expression of pathogenic fusion proteins. Leukemia features a particular high frequency of fusion oncogenes^1^ and the functional investigation of leukemia-associated fusion proteins has provided invaluable insights into the molecular mechanisms of cancer development^2^.

The protein complexes around fusion oncoproteins play defining roles in shaping oncogenic gene expression patterns^3^. Thus, the investigation of protein interactions is critical for uncovering actionable targets to devise more effective and better targeted cancer therapies. Several studies have used affinity-purification coupled to mass spectrometry (AP-MS) to identify novel key effectors of fusion-protein-related leukemia^4-6^. Yet, while the most common leukemia fusion proteins have been extensively characterized, functional understanding of the many rare fusions which affect a significant number of patients and have limited treatment options is lacking.

The N-terminal part of the Nucleoporin 98 (*NUP98*) gene (N-NUP98) is fused to over 30 different C-terminal partner loci in acute myeloid leukemia (AML)^7^. While *NUP98* rearrangements are rare (~2% of all AML)^7,8^, they are more frequent in pediatric AML and are associated with a particularly bad prognosis^9^.

The endogenous NUP98 protein is part of the nuclear pore complex (NPC), which mediates bidirectional transport of macromolecules between nucleus and cytoplasm10,^11^. NUP98 is part of the group of FG Nucleoporins, which contain intrinsically disordered regions (IDRs) consisting of repeats of either Phe-Gly (FG), or Gly-Leu-Phe-Gly (GLFG) amino acid residues. GLFG repeats can serve as binding sites for RNA-binding proteins, like the mRNA export factor RAE1, thus mediating trafficking of RNA molecules through the NPC^7,12^. While the majority of mature NUP98 protein is directly recruited to the NPC, NUP98 was also found to associate with the anaphase promoting complex (APC)^13^ and to interact with chromatin, where it actively regulates gene expression in an NPC-independent fashion^14,15^.

C-terminal fusion partners of N-NUP98 in AML are enriched for proteins with roles in transcriptional control and epigenetics. Several studies have addressed the functional contribution of different protein modules present in NUP98-fusions to leukemogenesis. For instance, the plant homeodomain (PHD) in NUP98-PHF23^16^ and the RNA helicase motif in NUP98-DDX10 ^17^ were required for leukemogenesis, demonstrating critical roles for the C-terminal fusion partner. Conversely, deletion of the N-terminal NUP98-moiety in NUP98-NSD1 also prevented myeloid progenitor immortalization and high HOX gene expression, highlighting the importance of the conserved IDR domain for oncogenic transformation of NUP98-fusion proteins^18^. Yet, the exact molecular mechanisms of NUP98-fusion protein-induced leukemogenesis remain poorly understood, and it is not clear which of the oncogenic properties of NUP98-fusion proteins depend on molecular functions of endogenous NUP98 are governed by novel functions that are mediated by the fusion partner^19^. We hypothesized that NUP98-fusion proteins interact with specific networks of cellular proteins, and that oncogenic functions of NUP98-fusion proteins are hard-wired in their protein interactomes.

Here we show by AP-MS-based interactome analysis that NUP98-fusion proteins do not act in the context of the NPC. Consistent with the specific nuclear localization pattern of NUP98-fusion proteins, the core NUP98-fusion interactome is highly enriched for proteins with known roles in liquid-liquid phase separation (LLPS) and the formation of biomolecular condensates. To investigate global changes in biomolecular condensation we established *biotinylated isoxazole-mediated condensome mass spectrometry* (biCon-MS), a sensitive method to globally characterize the dose-dependent potential of IDR-containing cellular proteins to form precipitates in the presence of the chemical biotinylated-isoxazole (b-isox). We show that biCon-MS greatly expands the cellular catalogue of proteins involved in LLPS beyond known protein complexes that act in biomolecular condensates. Furthermore, biCon-MS revealed that NUP98-fusion protein expression specifically alters the composition of cellular condensomes, indicating that NUP98-fusion-driven oncogenesis involves altered biomolecular condensation. In fact, an artificial FG-repeat-containing IDR-fragment fused to the C-terminal fusion partner KDM5A was capable of inducing leukemia-associated gene expression. Our data show that the biophysical properties of oncogenic fusion protein partners have the potential to specifically alter cellular biomolecular condensation to drive cancer-specific gene expression programs.

## Results

### NUP98-KDM5A does not operate in the context of the nuclear pore complex

As the contribution of physiological functions of endogenous NUP98 to NUP98-fusion-dependent oncogenesis is unclear, we decided to compare global protein-protein interactions of endogenous NUP98 with those of a NUP98-KDM5A fusion protein. A Strep-HA-tagged variant of NUP98-KDM5A (Figure 1a) was stably expressed in the human leukemia cell line HL-60. In line with previous results^20^, NUP98-KDM5A did not co-localize with the nuclear membrane but was present in intra-nuclear speckles (Figure 1b). To enable unambiguous annotation of protein complexes to endogenous NUP98 vs. NUP98-KDM5A, we purified endogenous NUP98-protein complexes from HL-60 cells using a highly specific anti-NUP98 antibody, while the NUP98-KDM5A interactome was isolated via the Strep-tag present in the fusion protein (Figure 1c). Enrichment of baits was confirmed by Western blot and functional purification of protein complexes was shown by co-precipitation of the known NUP98-binding partner RAE1 ^21^ (Figure S1a). In line with its altered localization, NUP98-KDM5A did not coprecipitate with NUP98 (Figure S1a).

**Figure 1.**
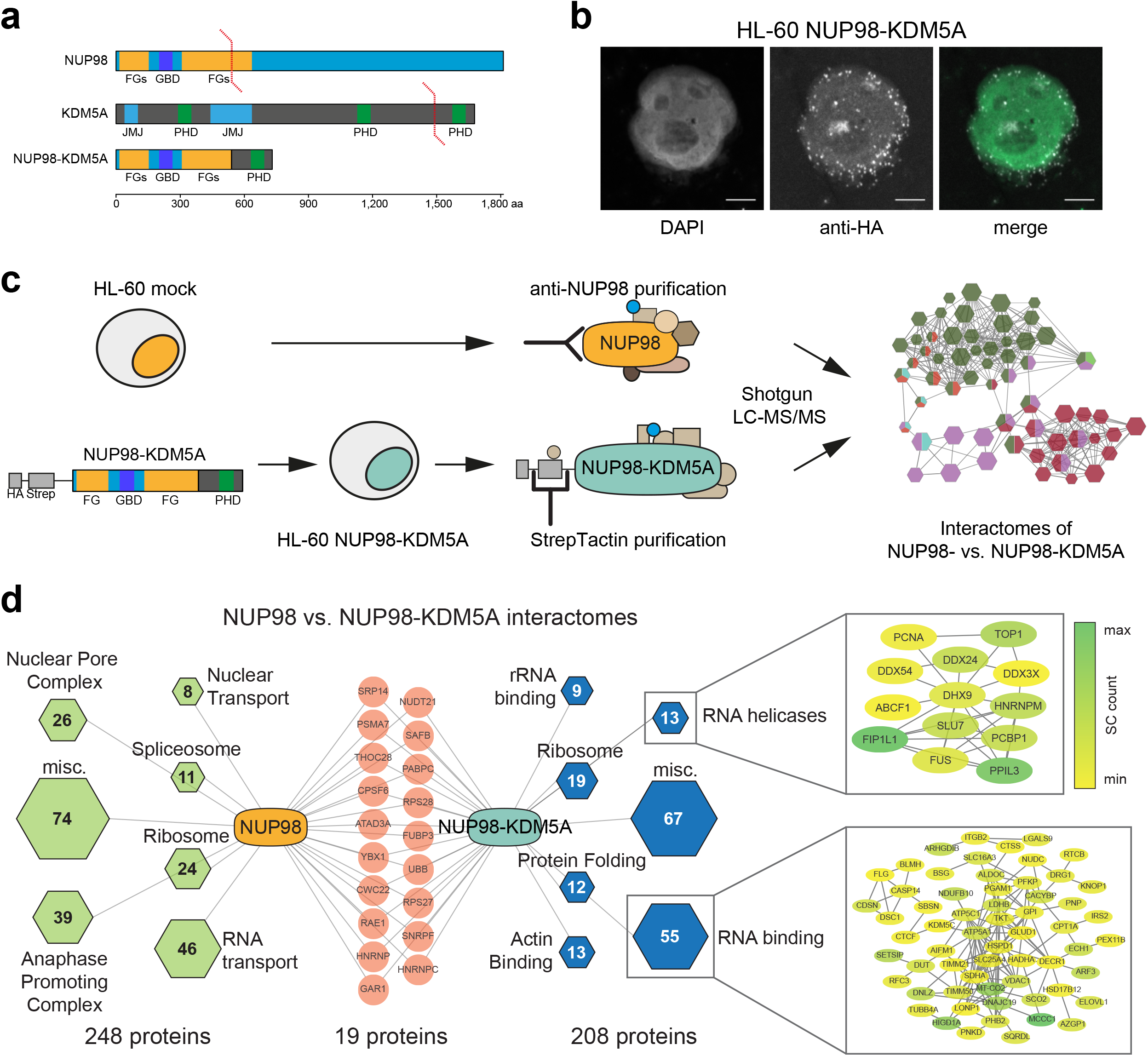
NUP98-KDM5A does not operate in the context of the nuclear pore complex. (a) Schematic illustration of endogenous NUP98 and KDM5A proteins, as well as the oncogenic fusion protein NUP98-KDM5A. FG – Phenylalanin/Glycin, GBD – Gle2 binding sequence domain, JMJ – Jumonji domain, PHD – plant homeodomain. (b) Confocal microscopy images of NUP98-KDM5A-expressing HL-60 cells stained with DAPI and anti-HA antibody for exogenous fusion proteins. Scale bar: 5 μm. (c) Schematic representation of transduction of HL-60 cells with NUP98-KDM5A (left). Pulldown of endogenous NUP98-complexes in HL-60 mock cells with a NUP98-specific antibody and Strep-Tactin purification of tagged NUP98-KDM5A-complexes in transduced HL-60 cells (middle). Data acquisition of purified protein complexes using liquid chromatography coupled to mass spectrometry (LC-MS/MS) and subsequent data analysis (right). (d) Interactome analysis of NUP98 and NUP98-KDM5A. Individual protein complexes within the interactomes were obtained by K-means clustering, assigned using GO Biological Processes and illustrated as hexagons (green for NUP98- and blue for NUP98-KDM5A-complexes). Size of the hexagons and numbers represent the identified proteins associated to respective subcomplexes. Red nodes show shared proteins between the interactomes. Detailed STRING networks are shown for subcomplexes of “RNA helicases” and “RNA binding” and individual proteins are highlighted according to abundance in MS acquisition.

Purified protein complexes were analyzed by liquid chromatography coupled to mass spectrometry (LC-MS/MS) using a one-dimensional gel-free approach, identifying 315 and 390 proteins in anti-NUP98- and in STREP-tag-mediated purifications of NUP98-KDM5A, respectively. After stringent filtering for background and non-specific interactions ^22^, the NUP98 interactome featured 267 proteins (Table S1), and the NUP98-KDM5A interactome consisted of 227 proteins. The interactome of endogenous NUP98 recapitulated protein complexes that were previously reported ^15^ to interact with NUP98 with high affinity, including the NPC (26 proteins) ^23^, the APC (39 proteins) ^24^ and factors involved in nuclear transport (8 proteins) (Figure 1d, S1B). In contrast, NUP98-KDM5A mainly co-purified with distinct sets of RNA binding protein complexes, including RNA helicases (13 proteins) and RNA binding factors (55 proteins) (Figure 1d, S1b). Most importantly, only 19 of 497 proteins interacted with both endogenous NUP98 and NUP98-KDM5A. Among those was RAE1, which was reported to bind the N-terminus of NUP98 ^25^ (Figure 1d). These data suggest that despite sharing the same N-terminal sequence, NUP98- and NUP98-KDM5A operate in largely nonoverlapping cellular contexts, and that NUP98-KDM5A is not co-localizing with the NPC.

### Functional proteomic identification of conserved interactors of diverse NUP98-fusion proteins

Given the structural heterogeneity of the >30 NUP98-fusion partners found in AML, it was unclear if potential effector mechanisms that are critical for NUP98-fusion dependent leukemogenesis might converge on a shared set of conserved interaction partners. To characterize the conserved interactome of NUP98-fusion proteins, we chose four molecularly diverse oncoproteins in addition to NUP98-KDM5A. We based our selection of fusion proteins both on their abundance in the AML patient population, but also on functional diversity of endogenous partner proteins. The most frequent fusion partners of NUP98 with KDM5A (JARID1A), a histone 3 lysine 4 (H3K4) di- and tri-demethylase ^26,27^, and NSD1, a histone methyltransferase (HMT) for H3K36 and H4K20 ^28^. NUP98-HOXA9 was chosen to represent the recurrent fusions of NUP98 to members of the *HOX* gene cluster. HOXA9 is a transcription factor that is highly expressed in hematopoietic stem/progenitor cells ^29^. Finally, fusions of NUP98 with the RNA helicase DDX10 and the transcriptional co-activator PSIP1 (LEDGF) were also included in this study. Strep-HA tagged variants of all fusion proteins were expressed in HL-60 cells and transgene expression was verified by Western blotting (Figure 2a). Confocal microscopy of transduced cells revealed that all NUP98-fusion proteins showed comparable patterns of speckled localization across the cell nucleus (Figure 1b, Figure 2b). Protein complexes nucleated by different NUP98-fusion proteins were purified from lysates of HL-60 cells and their composition was analyzed by LC-MS/MS (Figure S2a). Analysis of AP-MS data yielded 616 proteins engaging in over 6000 interactions. After subtracting proteins from lysates of non-transduced HL-60 cells and filtering for non-specific interactions, 501 differentially enriched proteins were retained as high-confidence interactors of NUP98-fusion proteins (Table S2). Each individual NUP98-fusion protein had between 19 and 61 exclusive interactions. In contrast, the majority of proteins in the network interacted with more than one NUP98-fusion protein (Figure 2c). Overall, 157 proteins were found in three or more NUP98-fusion interactomes, and a conserved set of 27 proteins was present in all five NUP98-fusion protein complexes (Figure 2c). This conservation of interaction partners indicates significant functional overlap among different NUP98-fusions, pointing to the presence of similar molecular mechanisms. The 157 conserved NUP98-fusion interactors were enriched in protein complexes involved in RNA splicing, ribosome biogenesis and transcriptional control (Figure 2d). Further analysis using Gene Ontology (GO) annotation confirmed a significant enrichment for DNA- and RNA-related processes, including mRNA processing and transcription, which is in line with initial results from the NUP98-KDM5A interactome (Figure 2e).

**Figure 2.**
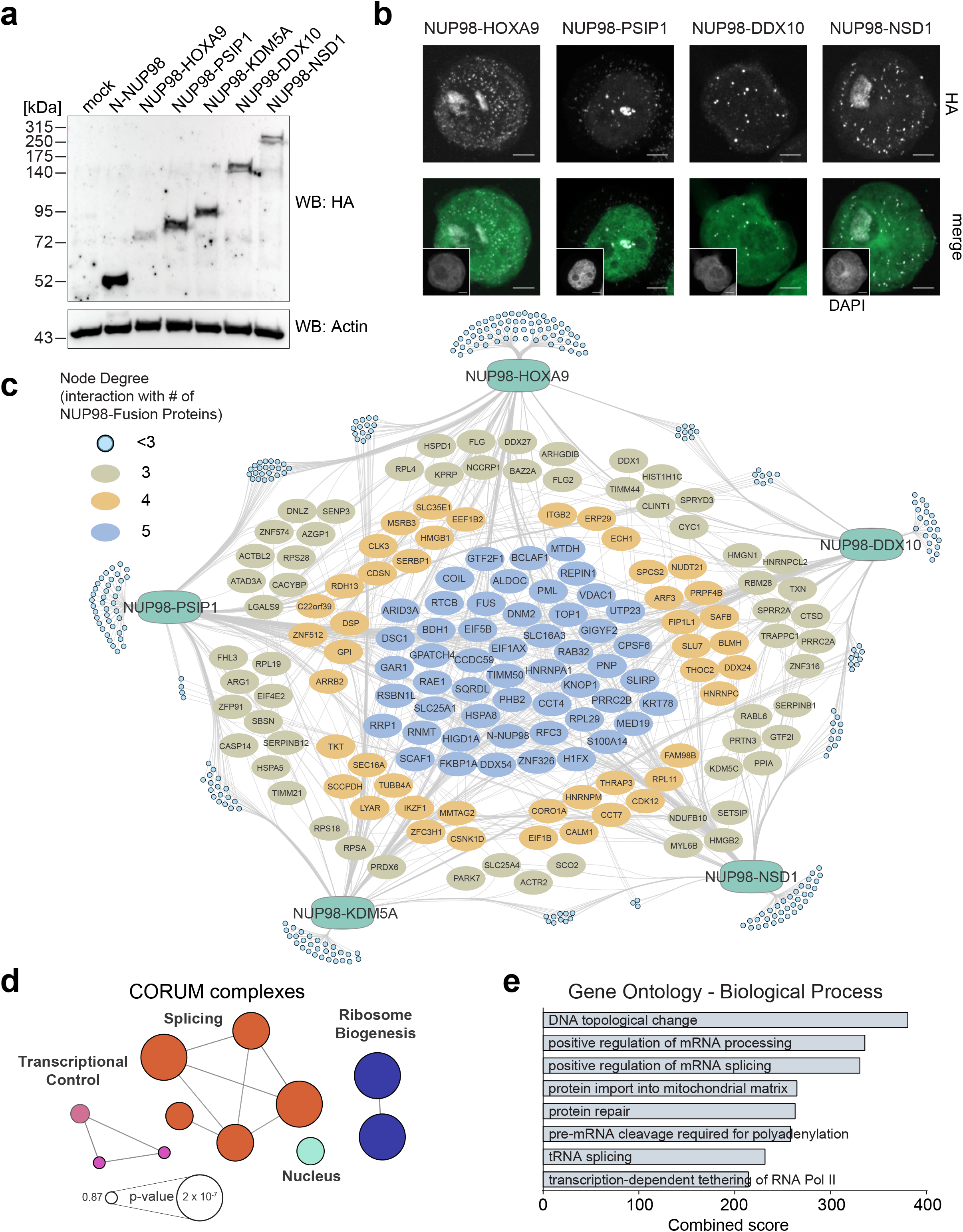
Functional proteomic identification of conserved interactors of diverse NUP98-fusion proteins. (a) Western blot analysis of mock-transfected, N-NUP98- or NUP98-fusion protein-expressing HEK293T cells. Exogenous proteins were detected with anti-HA antibodies and β-Actin was used as a loading control. (b) Confocal microscopy images of HL-60 cells expressing different NUP98-fusion proteins stained with DAPI and anti-HA antibodies for exogenous fusion proteins. Scale bar: 5 μm. (c) Interactome analysis of five NUP98-fusion proteins. Light blue nodes represent proteins interacting with less than three baits. Proteins that interact with three or more baits are illustrated as ovals. MS analysis was performed in 4 biological replicates. (d) CORUM analysis of core interactors. Complexes are illustrated as network highlighting most significant terms. Generated with ClueGo v3.5.4. (e) GO analysis for biological processes was performed for 157 interactors shared by at least three of five NUP98-fusion proteins. GO terms were ranked by combined score using Enrichr.

Notably, this proteomic survey of different NUP98-fusion proteins does not show colocalization with the nuclear pore complex. The conserved core interactome of NUP98-fusion proteins is enriched in factors regulating mRNA metabolism and transcription, pointing to an involvement of NUP98-fusion proteins in these processes.

### The NUP98-fusion protein interactome is enriched for proteins with roles in biomolecular condensation

Biomolecular condensates are membrane-less structures that govern biological processes through the dynamic compartmentalization of macromolecules^30^. Their formation can be driven by multivalent, low-affinity interactions between IDR-containing proteins, which are able to undergo LLPS^31^. Nuclear biomolecular condensates mediate important regulatory roles in transcription, splicing and chromatin organization^32^. Several well-described factors with roles in biomolecular condensation were present in the conserved NUP98-fusion protein interactome, including the RNA-binding protein FUS^33^, heterogeneous nuclear ribonucleoprotein A1 (HNRNPA1)^31^ and H/ACA ribonucleoprotein complex subunit 1 (GAR1)^34^.

To evaluate a potential enrichment of IDR-containing proteins involved in biomolecular condensation among the NUP98-fusion protein interactome, we used an algorithm that was designed to predict phase separation properties of proteins in an unbiased fashion. The PScore classifies the potential of proteins to self-associate based on the abundance of pi-orbital containing amino-acid residues, as pi-pi interactions are important features in phase separation^35^. Binning of NUP98-fusion-interacting proteins based on their PScores (STAR methods) revealed a significant overrepresentation of LLPS-prone proteins (Pscore >4) within functional categories that are enriched in the NUP98-fusion interactome, including RNA metabolism, RNA binding, RNA processing and transcription (Figure 3a). In line with this, the mean PScore of the individual 157 core interactors was significantly higher than a size-matched list of the human proteome that was obtained by random subsampling (Figure 3b). In contrast, the mean PScore of proteins assigned to the GO category “Nuclear Membrane” did not significantly divert from the human proteome (Figure S3a), and the subsampling process did not alter the global distribution of PScores (Figure S3b).

**Figure 3.**
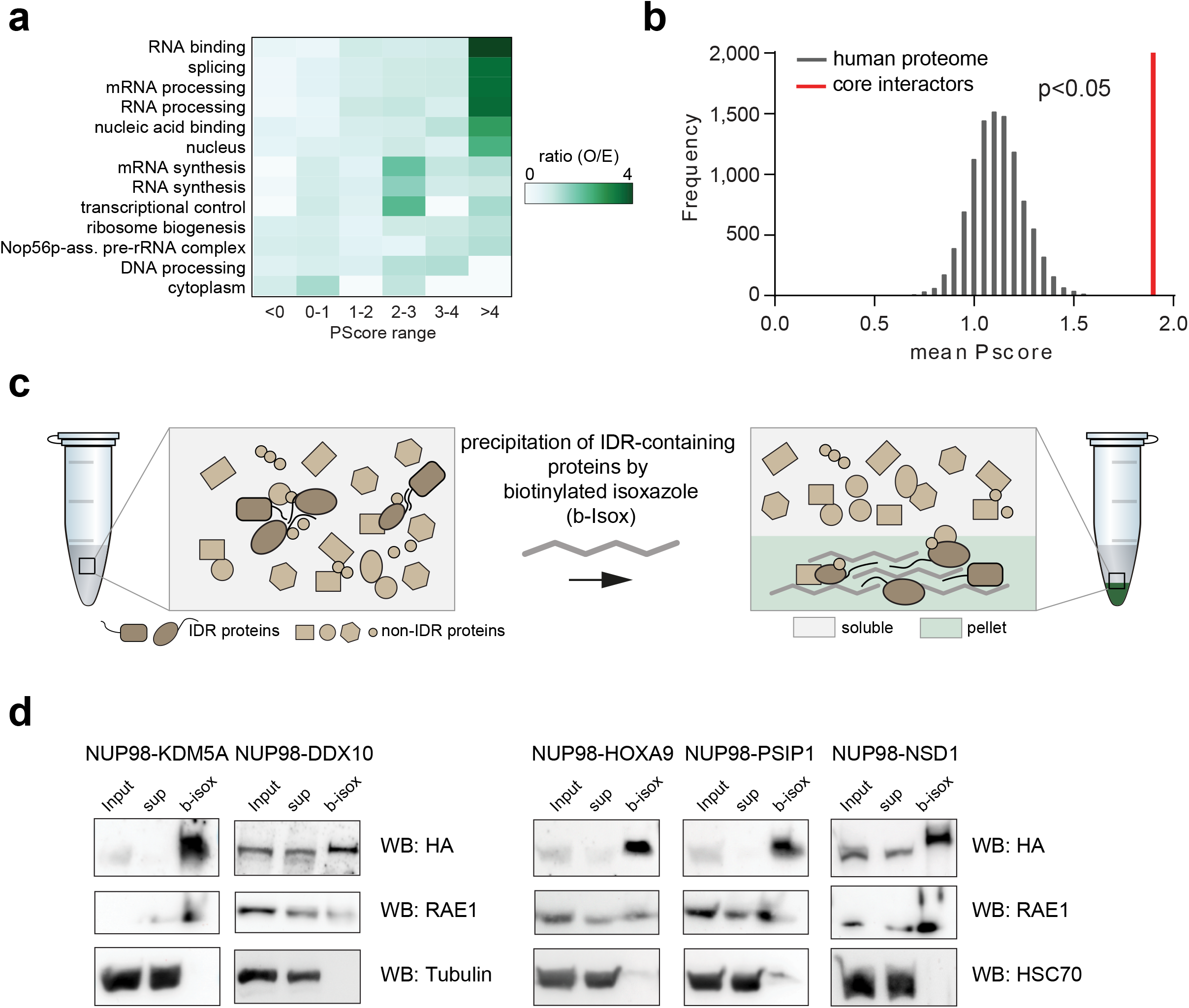
The NUP98-fusion protein interactome is enriched for proteins with roles in biomolecular condensation. (a) Observed/expected (O/E) ratios of binned PScores for proteins of significantly enriched Gene Ontologies were calculated by the observed PScore distribution of the core interactors compared to the expected distribution of the proteome. Significant GO terms were ranked by O/E ratio of the PScore >4 bin. (b) The mean PScore for NUP98-fusion core interactors was compared to a list of randomly subsampled lists from the human proteome of the same size. The P-value was calculated using the Kolmogorov-Smirnov test. (c) Schematic illustration of the b-isox-mediated precipitation assay. IDR-containing proteins form beta sheets upon b-isox treatment and can be collected by centrifugation. (d) Western blot analysis of HL-60 cell lysates expressing different NUP98-fusion proteins treated with 100 μM b-isox. Fusion proteins were detected with anti-HA antibodies, RAE1 with anti-RAE1 antibody and housekeeping with anti-HSC70 or anti-β-Actin. Sup, supernatant.

Given the enrichment of LLPS-prone factors among the core NUP98-fusion interactome we next aimed to investigate the biophysical properties of NUP98-fusion proteins with regard to biomolecular condensation. The chemical substance biotinylated isoxazole enables precipitation of IDR-containing proteins via the formation of microcrystals in solution (Figure 3c)^36,37^. All NUP98-fusion proteins precipitated efficiently upon incubation with 100 μM b-isox (Figure 3d). This effect was likely mediated by the IDR domain in the NUP98-N-terminus, which was previously shown to undergo LLPS *in vitro* ^38^. Despite the absence of IDR domains, the known NUP98 interactor RAE1 was efficiently co-precipitated with NUP98-fusion proteins in this assay, while the highly structured heat shock cognate 71 kDa protein (HSC70), which does not interact with NUP98-fusion proteins, was insensitive to b-isox precipitation (Figure 3d).

Our data show that the interactome of NUP98-fusion proteins is enriched for proteins with biophysical properties that are predictive of biomolecular condensation via LLPS. Consistent with their speckled nuclear localization, we find that NUP98-fusion proteins and interacting proteins are efficiently precipitated by b-isox, indicating that this assay allows investigation of the condensation behavior of protein assemblies within complex cellular lysates. Taken together, our results indicate that NUP98-fusion proteins and their interactomes are involved in biomolecular condensation in AML cells.

### biCon-MS globally charts the cellular condensome

Given the potential of the b-isox precipitation assay to capture protein complexes that are prone to biomolecular condensation, we attempted to leverage this effect to analyze the entirety of cellular proteins that are sensitive to b-isox in an unbiased fashion using mass spectrometry. We developed an experimental setup that allows the characterization of subsets of the cellular proteome that are dynamically integrated in b-isox-precipitates in a dose-dependent manner (Figure 4a, S4a,b). This optimized method, which we termed *biotinylated isoxazole-mediated condensome mass spectrometry* (biCon-MS) efficiently reduces background noise caused by non-specific precipitation and increase the likelihood of true positive hits by eliminating proteins that do not display dose-dependent enrichment, as exemplified by Western blot analysis for the NUP98-interaction partner RAE1 (Figure 4b). biCon-MS analysis of HL-60 lysates identified 931 proteins that exhibited dose-dependent precipitation behavior (Table S3), while 209 proteins were significantly enriched in 33 μM precipitates compared to 11 μM. Well-described factors implicated in biomolecular condensates such as FUS, TAF15 and MED12 were retrieved in a dose-dependent fashion (Figure 4c). Pscore analysis indicated that these 209 proteins were highly enriched for proteins capable of LLPS (Figure 4d). Functional annotation of proteins showing dosedependent b-isox precipitation behavior resulted in a comprehensive network that was representative of diverse cellular physiology associated with biomolecular condensation including cellular signaling and metabolism but also RNP assembly, chromatin organization and gene expression (Figure 4e). Further clustering, based on interactions reported in the STRING database, of proteins in the most significantly enriched hub (“gene expression”) generated a network of 81 proteins with 496 interactions. It consisted of different complexes that play critical roles in transcriptional regulation (transcription factors GFI1, FLI1 and RUNX2), transcriptional initiation (components of the Mediator complex, the SWI/SNF complex and RNA Polymerase II), but also proteins involved in RNA binding and translation, such as TAF15 or HNRNPA2B1 (Figure 4e). Importantly, many of these factors have been previously implicated in LLPS, which is consistent with their high PScores (Figure 4e), providing further evidence of this algorithm’s suitability as an unbiased predictor of the capability of individual proteins to participate in biomolecular condensation.

**Figure 4.**
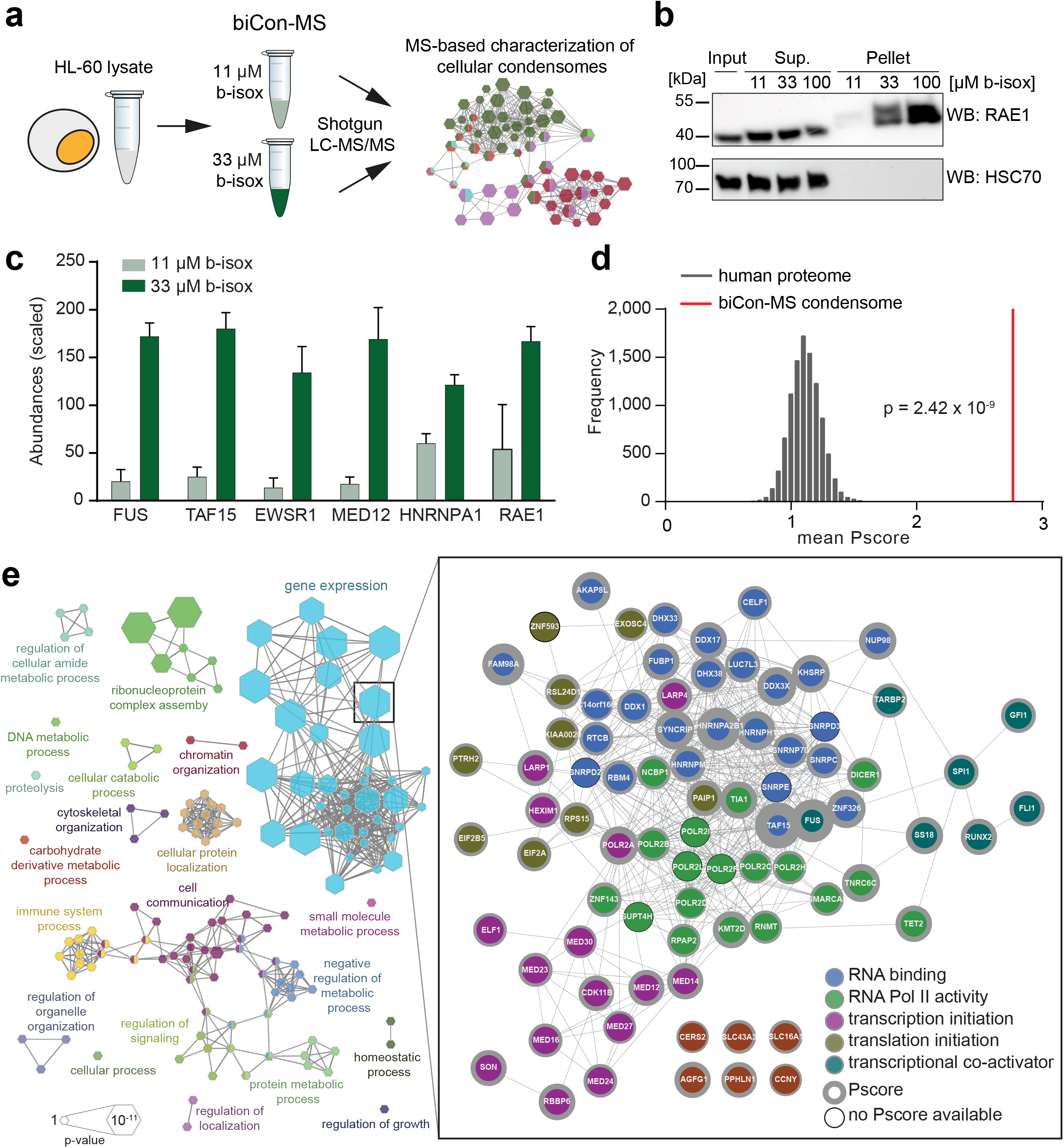
biCon-MS globally charts the cellular condensome. (a) Schematic illustration of the biCon-MS approach (b) Western blot analysis of HL-60 cell lysates treated with 11 μM, 33 μM or 100 μM b-isox. Dose-dependent precipitation was investigated for RAE1 and HSC70. (c) Examples of proteins identified by biCon-MS in HL-60 cells. Normalized and scaled LFQ-based protein abundance as calculated by Proteome Discoverer. (d) Mean PScore for proteins enriched (p-value < 0.05, log2(fc) > 0.5) in 33 μM b-isox precipitates compared to 11 μM b-isox precipitates was compared to a list of randomly subsampled lists of the human proteome with the same size. P-value was calculated using the Kolmogorov-Smirnov test. (e) Gene Ontology analysis of proteins enriched (p-value < 0.05, log2(fc) > 0.5) in 33 μM b-isox precipitates compared to 11 μM b-isox precipitates. Edges connect terms with overlapping protein lists. The protein list of the most significant GO term (“gene expression”) is represented by interactions from STRING. The network of annotated interactions was clustered using Reactome FI in Cytoscape and colored accordingly. Gray border thickness indicates PScores for individual proteins. Proteins without any annotated PScore are indicated by black borders. Size of the hexagons scale with significance.

Together, these data show that biCon-MS analysis of cellular lysates represents a powerful approach to investigate the global composition of the cellular condensome, providing an unbiased and comprehensive map of cellular proteins that are involved in biomolecular condensation.

### Expression of NUP98-fusion proteins dynamically alters the cellular condensome

As we found that NUP98-fusion proteins specifically interact with proteins capable of LLPS, we next aimed to investigate dynamic changes in the cellular condensome that result from the expression of oncogenic NUP98-KDM5A or NUP98-NSD1 fusion proteins by biCon-MS (Figure 5a). In addition to proteins that consistently present in biCon-MS, such as FUS, TAF15, EWSR1 (Figure S4c and S4d), we found a dose-dependent increase of NUP98-fusion proteins, confirming that NUP98-fusion proteins are involved in biomolecular condensation (Figure 5b). Comparison of global condensomes in NUP98-fusion protein expressing- vs. mock HL-60 cells based on stringent statistical filtering revealed extensive reorganization of the cellular condensome upon oncogene expression (Figure 5c, Table S3). 145 and 240 proteins were significantly depleted from cellular condensomes upon expression of NUP98-KDM5A and NUP98-NSD1, respectively. Conversely, 64 and 70 proteins were significantly enriched in condensomes of NUP98-KDM5A- and NUP98-NSD1-expressing cells (Figure S4e,f). In line with the hypothesis that NUP98-fusion proteins share molecular mechanisms leading to oncogenic transformation, the majority of differentially enriched proteins in fusion protein condensomes were shared between NUP98-KDM5A and NUP98-NSD1 (Figure 5d). Network analysis of all proteins enriched in condensomes upon NUP98-fusion protein expression uncovered key complexes of transcriptional activation and chromatin organization (Figure 5e). For instance, different subunits of the Mediator complexes such as MED31 and MED8 were exclusively found in both NUP98-fusion protein-condensomes (Figure 5e). In contrast, MED15 was significantly depleted from condensomes upon expression of NUP98-KDM5A and NUP98-NSD1 when compared to parental HL-60 cells (Figure 5f). In addition, important factors in leukemia development, such as RUNX2 and TET2 were also enriched in NUP98-fusion dependent condensomes. (Figure 5f).

**Figure 5.**
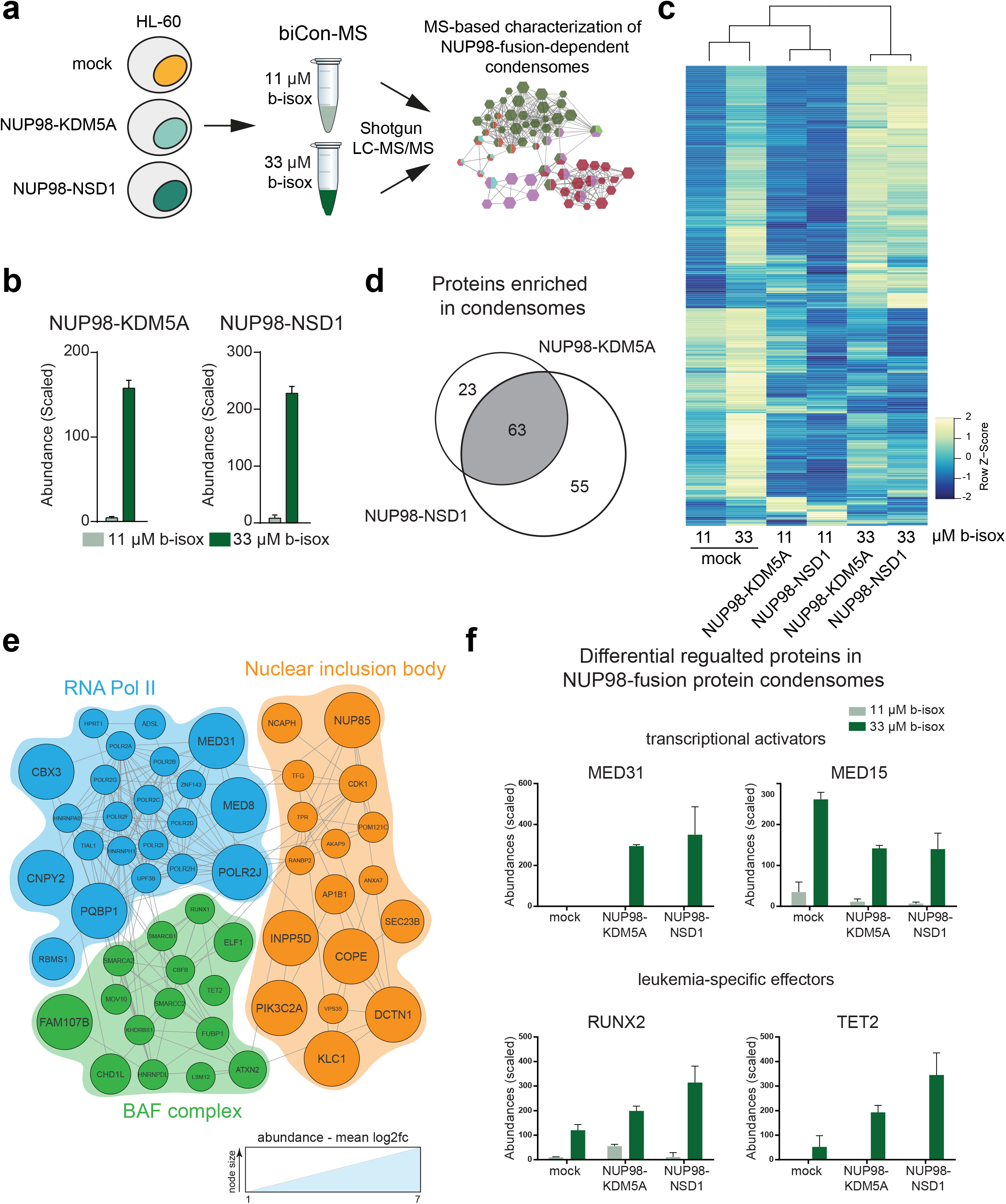
Expression of NUP98-fusion proteins dynamically alter the cellular condensome. (a) Schematic illustration of biCon-MS for HL-60 cells expressing NUP98-KDM5A or NUP98-NSD1. (b) Scaled abundance for Strep-HA-tag-derived peptides identified by biCon-MS in lysates of NUP98-fusion protein-expressing HL-60 cells. (c) Heatmap of all proteins that were precipitated more abundantly in mock HL60 cells with 33 μM b-isox than in mock HL60 cells with 11 μM HL60 cells. Rows and columns were clustered using Pearson correlation as distance measure and ward.D clustering. Each row represents Z-scores of scaled mean abundances for individual proteins for each condition. (d) Venn diagram of significantly enriched proteins in condensomes of NUP98-KDM5A- and NUP98-NSD1-expressing cells when compared to mock HL-60 cells. (e) Reactome FI clustering of all proteins that precipitated by b-isox in a dose-dependent manner upon expression of NUP98-KDM5A and NUP98-NSD1 as compared to mock HL60 cells. Proteins representative for the three most significant protein complexes are shown in groups and connected via annotated STRING data base interactions. Size of nodes represent mean log2(fc). (f) Scaled abundance of significantly enriched proteins in both NUP98-fusion protein condensomes as identified by biCon-MS.

In summary, these data show that NUP98-fusion proteins cause extensive restructuring of the cellular condesome, supporting the hypothesis that NUP98-fusion proteins are able to specifically reconstitute biomolecular condensates.

### An artificial IDR-containing KDM5A fusion protein is sufficient to drive leukemia-associated gene expression

While our data show that NUP98-fusion proteins are involved in biomolecular condensation, it was not clear if this function plays a role in NUP98-fusion-mediated transcriptional reprogramming of myeloid progenitor cells to induce leukemia. The IDR-containing N-terminus of NUP98 is essential for the formation of biomolecular condensates *in vitro* ^38^. To test whether the biophysical properties of an FG-repeat-containing IDR fused to the KDM5A C-terminus is sufficient to induce leukemia-specific gene expression patterns, we designed an artificial peptide containing 39 FG-repeats and fused it to the C-terminus of KDM5A, resulting in an artificial FG-KDM5A fusion protein (artFG-KDM5A). As a control, the KDM5A C-terminus was fused to a sequence of 39 di-peptides of the nonpolar and aliphatic amino acid alanine (A) (artAA-KDM5A). artFG-KDM5A showed speckled nuclear localization (Figure 6b) that was similar to NUP98-KDM5A (Figure 1b), while artAA-KDM5A was distributed homogenously across the cell nucleus (Figure 6b). Constructs were introduced into mouse fetal liver stem cells and resulting changes in gene expression were compared to control and the effects of the original NUP98-KDM5A fusion protein by RNA-seq (Figure 6a). Principal component analysis of RNA-seq data revealed high similarity of global gene expression between artAA-KDM5A and wild-type (WT) fetal liver cells, while transcriptional profiles resulting from expression of NUP98-KDM5A and artFG-KDM5A were significantly distinct (Figure S5a). Differential gene expression analysis revealed that the pattern of deregulated genes in NUP98-KDM5A- and artFG-KDM5A-expressing cells differed significantly from wild type cells. In contrast, only minor changes were observed upon expression of artAA-KDM5A (Figure 6c). Out of 987 genes whose expression was deregulated by NUP98-KDM5A, artFG-KDM5A expression caused consistent expression changes in 761 genes (78%) (Figure 6d). In line with this, KEGG pathway analysis of differentially expressed genes in NUP98-KDM5A- and artFG-KDM5A-expressing cells revealed a high overlap in affected pathways (Figure 6e). Strikingly, the 761 genes differentially regulated in both cell populations, were highly characteristic of gene expression patterns for hematopoietic malignancies (Figure S5b), highlighting the oncogenic transcriptional potential of the artFG-KDM5A fusion protein. Finally, we found that exogenous expression of the artFG-KDM5A fusion was able to induce high expression levels of the direct transcriptional targets of NUP98-fusion proteins that we have recently identified, including *Hoxa7, Mira* and *Cdk6*^39^ (Figure 6f).

**Figure 6.**
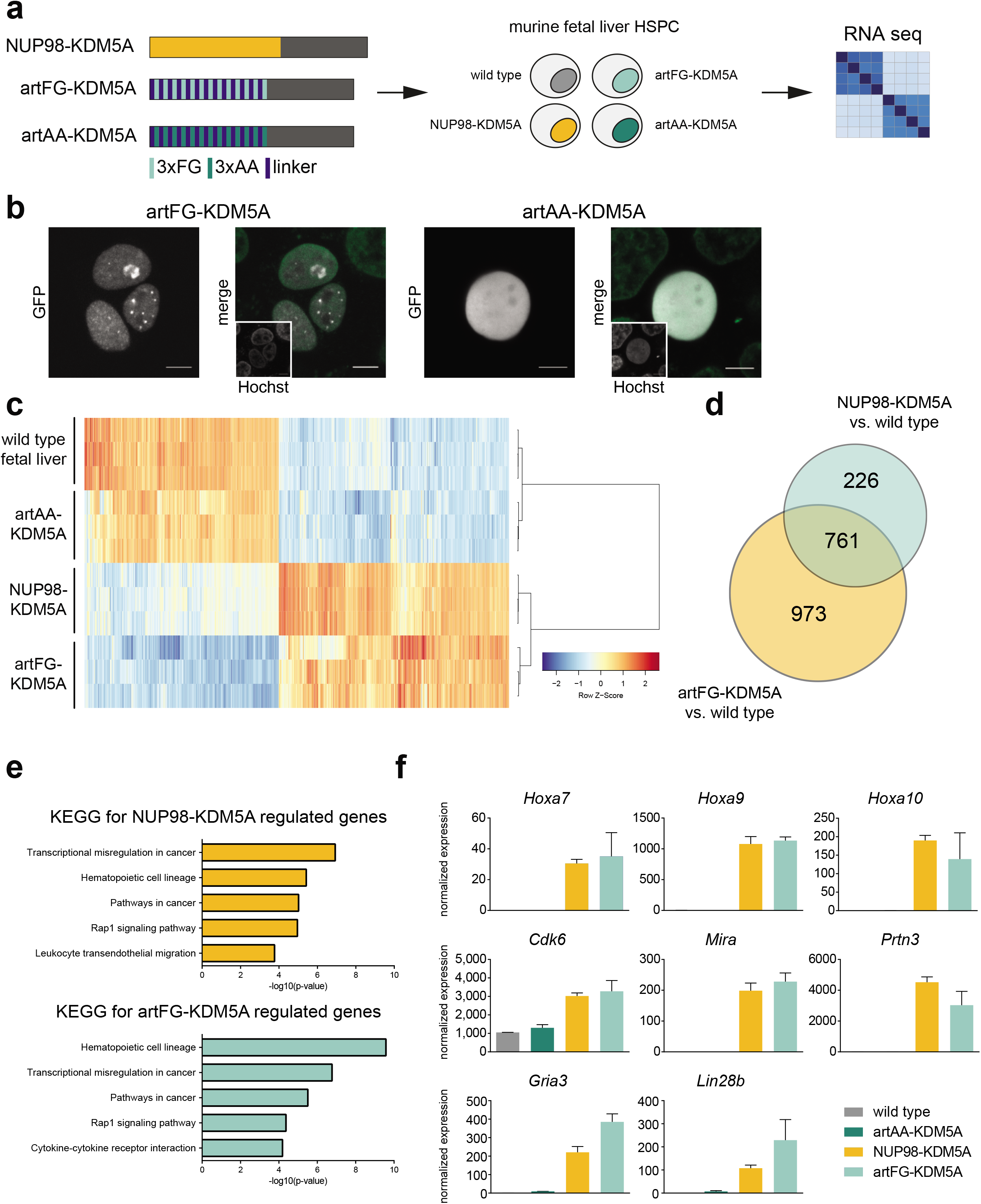
Distinct IDR-containing fusion proteins transform fetal liver cells *ex vivo*. (a) Schematic illustration of the experimental approach. (b) Live cell imaging of HEK293-T cells expressing GFP-tagged variants of artFG-KDM5A and artAA-KDM5A. Fusion proteins are shown in white and Hoechst staining is shown in green. Scale bar 5 μm. (c) Heatmap of significantly de-regulated genes in NUP98-KDM5A- and artFG-KDM5A-expressing cells compared to control fetal liver stem cells. Rows and columns were clustered using Pearson correlation as distance measure and ward.D clustering. Each row represents Z-scores of scaled expression levels for each replicate. Only genes with p-value < 0.01 are shown. (d) Venn diagram of differentially regulated genes in NUP98-KDM5A and artFG-KDM5A expressing cells compared to mock. P-value < 0.001 and log2 fold change < -2 or > 2. (e) KEGG pathway analysis for differentially regulated genes of NUP98-KDM5A (top) and artFG-KDM5A (bottom) expressing fetal liver cells. (f) Normalized expression of direct NUP98-fusion protein targets.

Taken together, these results show that the biophysical properties of FG-repeat-containing IDRs are sufficient to phenocopy the gene-regulatory potential of the NUP98 N-terminus in the context of clinically relevant NUP98-fusion oncoproteins.

## Discussion

In this study, we show that structurally different NUP98-fusion proteins localize to biomolecular condensates and that the structural determinants guiding biomolecular condensation as encoded in the NUP98 N-terminus are sufficient to evoke leukemia-specific gene expression in the context of oncogenic fusion proteins. Thus, we propose that alteration of biomolecular condensation mediated by IDR-containing oncogenic fusion proteins represents a novel mechanism of oncogenic transformation.

Our results provide the first comprehensive AP-MS-based interactome analysis of NUP98-fusion proteins vs. endogenous NUP98 in human cells. In line with previous reports, our data show that NUP98 is recruited to the nuclear pore complex^23^ and the anaphase promoting complex^24^. Furthermore, we confirm its important role in RNA transport via interaction with RAE1^25^ and the ATP-dependent RNA helicase A (DHX9)^15^. Out of 247 identified NUP98 interactors only 46 have not been annotated as part of the NUP98 protein complex, highlighting the high purity of the NUP98 interactome presented here. AP-MS-based comparison of the interactomes of endogenous NUP98 vs. a NUP98-KDM5A fusion protein revealed surprisingly little overlap. While most overlapping proteins (e.g. HNRNPA1, YBX1, NUDT21) are part of the RNA transport machinery, they might interact with endogenous NUP98 or NUP98-fusion proteins in distinct contexts. Extended interactome analysis of five structurally diverse yet clinically relevant NUP98-fusion proteins further demonstrated their association with protein complexes involved in RNA metabolism. A core set of 157 proteins represents the conserved core interactome of NUP98-fusion proteins. Functional annotation of this core NUP98-fusion interactome revealed high enrichment for proteins involved in the control of gene expression. These results support our hypothesis that structurally distinct NUP98-fusion proteins share highly specialized protein environments that are critical for their oncogenicity. Conversely, proteins that uniquely interact with individual fusion proteins might as well be critical for the oncogenic function of NUP98-fusion proteins. Strikingly, the NUP98-fusion core interactome was enriched in factors that were previously implicated in the formation of biomolecular condensates, such as FUS^33^, HNRNPA1^31^ and GAR1^34^. Biomolecular condensates are membraneless subcompartments within cells that serve to efficiently compartmentalize macromolecules^40^, such as stress granules^31^, Cajal bodies^41^ or paraspeckles^40^. Their formation is often governed by IDR-containing proteins^31^, which are capable of undergoing LLPS. Recent studies have identified a critical role for biomolecular condensation during transcriptional control, translation and protein transport^32,42-44^. For instance, IDR-containing transcription factors^43^ and coactivators undergo LLPS at superenhancers to establish subnuclear compartments that concentrate the transcriptional machinery to ensure efficient gene control^42^. Using the PScore, a computational approach to predict the propensity of proteins to undergo LLPS based on pi-pi interactions between and within proteins^35^, we found a significant enrichment of proteins capable to undergo LLPS among the 157 core interactors of NUP98-fusion proteins.

Several experimental approaches exist to characterize the localization and composition of biomolecular condensates. Confocal imaging is often used to directly visualize biomolecular condensates in cells^45^. We and others have observed that NUP98-fusion proteins show speckled localization in the nucleoplasm, but do not localize to the nuclear membrane^20,46,47^. This is consistent with our AP-MS data, supporting roles of NUP98-fusion proteins in RNA metabolism and gene expression in the context of biomolecular condensates. Alternatively, the b-isox-mediated precipitation assay can be used to assess the propensity of proteins to form aggregates in the context of complex cellular lysates^36,37^. The NUP98 N-terminus contains two large IDRs that consist of 38 di-amino acid repeats of phenylalanine-glycine (FG) and NUP98 is indeed able to form biomolecular condensates^38,48^. Consistent with the biochemical and biophysical properties of the NUP98 N-terminus, all NUP98-fusion proteins were highly susceptible to b-isox precipitation. This is in line with previous observations showing that NUP98 can localize to so-called “GLFG bodies” whose formation was dependent on NUP98 N-terminus ^49^.

To globally analyze the composition of biomolecular condensates in an unbiased manner and in a cellular context we developed biCon-MS. In this technique, cellular lysates are incubated with increasing concentrations of b-isox followed by MS-based identification of precipitated proteins. Beyond recovering the majority of proteins previously found to be sensitive to b-isox precipitation^36,37^, biCon-MS efficiently recovered several proteins with well-described roles in LLPS, such as the FET protein family (FUS/EWSR1/TAF15)^50^, 17 constituents of the mediator complex and 11 members of the RNA Pol II family^44^. Thus, biCon-MS greatly expands the cellular catalogue of proteins involved in biomolecular condensation. Functional annotation of these proteins clearly links biomolecular condensation to gene expression, mRNA transport and metabolic regulation, processes that were recently revealed to be critically dependent on LLPS^32,42–44,51,52^. Beyond that, we identified a large number of proteins that were not previously implicated in LLPS, including RNA helicases, epigenetic modulators and transcriptional co-activators.

biCon-MS analysis showed that expression of NUP98-KDM5A or NUP98-NSD1 fusion proteins caused substantial alterations in the composition of biomolecular condensates. In total, 134 proteins were significantly enriched in fusion-altered condensomes while 385 proteins were significantly depleted from condensomes upon NUP98 fusion protein expression. Core complexes involved in gene regulation were commonly enriched in condensomes in a NUP98-fusion dependent fashion. This includes factors that are essential for basic functions related to transcriptional control, such as POLR2A and MED31^53^, but also effects that have high relevance for leukemia-specific gene regulation, like RUNX2^54^ and TET2^55^. Specific, NUP98-dependent recruitment of these factors to biomolecular condensates involved in transcription might explain the unique effects NUP98-fusion proteins have on gene expression. Lastly, we were able to identify a large number of factors that have not yet been implicated in LLPS, and their potential involvement in pathogenic mechanisms is unclear. For example, adenylosuccinate lyase (ADSL) is a catalytic enzyme converting adenylosuccinate to adenosine monophosphate (AMP) and fumarate^56^. Thus, NUP98-fusion-specific alteration of enzyme function mediated by specific microenvironments that are present within biomolecular condensates might represent a novel aspect in targeting metabolic dependencies in cancer.

While the number of FG repeats had a strong influence on sub-nuclear localization of NUP98-fusion proteins^57^, and deletions in the NUP98 N-terminus affect the leukemogenicity of NUP98-HOX fusions^58^, it was not clear if the FG-repeats in the NUP98 N-terminus are a critical determinant of the oncogenicity of NUP98-fusion proteins. An artificially engineered IDR consisting of 13 triple FG-repeats fused to the C-terminal part of KDM5A was sufficient to phenocopy gene expression changes that are induced by the NUP98-KDM5A fusion, including induction of direct target genes of NUP98-fusion proteins. In contrast, a control construct containing di-alanine (AA)-repeats in place of the FG-repeat containing IDR was incapable of evoking NUP98-fusion specific gene expression changes. These results might provide a common molecular mechanism for how a family of >30 structurally diverse fusion proteins can cause a very similar disease. We postulate that the unique combination of IDR-containing protein sequences provided by the NUP98-terminus (mediating biomolecular condensation) together with protein modules involved in gene control (represented by the C-terminal fusion partners) dictates the oncogenic potential of NUP98-fusion proteins.

Investigation of the 368 factors that are involved in oncogenic fusion proteins as listed in the Catalogue of Somatic Mutations in Cancer (COSMIC) reveals a high enrichment of IDR-containing proteins, such as EWS, TAF15, NUP214 and FUS (Figure S5c). For instance, the EWS-FLI1 fusion protein, which is the driver oncogene in Ewing’s sarcoma, was shown to undergo phase transitions that are critical for gene activation^59^. Although the underlying molecular mechanisms might diverge, the capabilities of fusion proteins to undergo LLPS could indicate that alteration of biomolecular condensation is a fundamental principle of cancer development driven by oncogenic fusion proteins.

## Online Methods

### Constructs

cDNAs for different NUP98-fusion proteins were cloned into retroviral vectors enabling Tetracycline-inducible expression of Strep-HA-tagged-transgenes (pSIN-TREt-SH-gw-IRES-GFP-PGK-BlastR) using the Gateway technology. For constitutive transgene expression, cDNAs were cloned into pMSCV-SH-gw-PGK-BlastR-IRES-mCherry. Artificial fusion proteins (artFG-KDM5A and artAA-KDM5A) were cloned into constitutive expression vectors tagged to EmGFP.

### Cell culture

HL-60, HEK293-T and NIH-3T3 cells were obtained from the German Collection of Microorganisms and Cell Cultures GmbH (DSMZ) and Platinum-E (Plat-E) cells were purchased from Cell Biolabs, Inc. HL-60 cells were modified to stably express the ecotropic receptor and the reverse Tetracycline transactivator protein (rtTA3) (HL-60-RIEP) and were cultured in RPMI 1640 (Gibco) supplemented with 10% fetal bovine serum (FBS, Gibco), 2 mM L-Glutamine (Gibco) and 100 U/ml Penicillin-Streptomycin (Gibco). HEK-293T, NIH-3T3 and Platinum-E cells were maintained in DMEM (Gibco) supplemented with 10% FBS, 2mM L-Glutamine and 100 U/ml Penicillin-Streptomycin. All cell lines have been tested for mycoplasma contamination. All cytokines were purchased from Peprotech (Rocky Hill, NJ, USA).

HL60-RIEP cells were retrovirally transduced to stably express TREt-HA-Strep-NUP98-FP-IRES-eGFP-PGK-Blasti and cultivated in the presence of Puromycin (2 μg/ml) and Blasticidin (10 μg/ml). 1 μg/ml Doxycycline (Sigma-Aldrich) was added to induce cDNA transcription 24 hours prior to harvesting and transgene expression was monitored by flow cytometry for GFP.

### Fetal liver cell culture

Primary mouse fetal liver stem cells were isolated from fetal livers of C57BL/6N mice between day E13.5 and E14.5. Cells were cultured in DMEM/IMDM (50:50% vol/vol), supplemented with 10% FBS, 100 U/ml Penicillin-Streptomycin, 4 mM L-Glutamine and 50 uM beta-mercaptoethanol in the presence of murine stem cell factor (mSCF, 150 ng/ml), murine interleukin 3 (mIL-3, 10 ng/ml) and murine interleukin 6 (mIL-6, 10 ng/ml) (all PreproTech).

### Retroviral transduction

For retroviral transductions, Plat-E cells were transiently transfected with pGAG-POL and retroviral expression vectors using the Polyethylenimine (PEI) protocol^60^. Virus-containing supernatant was harvested, filtered (0.45 μm), and supplemented with polybrene (final concentration 10 μg/ml, Merck Millipore / TR-1003-G). Target cells were spinoculated at 800 g for 45 min on two successive days.

### Immunofluorescence analysis

Cells were spotted on glass histology slides using a Shandon CytospinTM Centrifuge II and air-dried. Spots were fixed with 4% Formaldehyde (Histofix, Roth, P087.6) for 10 minutes at 4 °C. Cells were permeabilized with 0.2% Triton X100 in PBS for 10 min at room temperature, followed by 1 hour incubation with primary antibody in 2% BSA/0.2% Triton X100 in PBS in a wet chamber. For detection of exogenous fusion proteins a mouse anti-HA (Santa Cruz, sc-7392, clone F-7, RRID: AB_627809) antibody was used. Samples were incubated with a secondary Alexa Fluor 568 f(ab’)2-goat anti-mouse antibody (Thermo Fisher Scientific, A-21237, RRID: AB_2535806) for 1 hour at room temperature in a dark wet chamber. DNA was stained with DAPI (4’,6-Diamidino-2-Phenylindole, Dilactate, Biolegend, 522801) and slides were mounted using Entellan^®^ New (Merck, 1079610100).

Images were acquired using a Zeiss SM 880 Airyscan - confocal laser scanning microscope with the Zeiss ZEN-black software. Post-processing of images was performed using ZEN-blue and ImageJ for brightness and contrast enhancement.

### Live cell imaging

Cells were transiently transfected with 250 ng plasmid DNA and imaged after 72 h on a Zeiss SM 880 confocal microscope. Nuclei were stained with 5 μg/ml Hoechst 33342 (Thermo Fisher Scientific, H1399) for 7 minutes. Post-processing of confocal z-slice images were accomplished using ZEN-blue software and ImageJ for contrast enhancement.

### Cell lysis for protein harvest

10-100×10^6^ cells were harvested and washed with PBS. Frozen cell pellets were resuspended in ice-cold AP-MS buffer (50 mM HEPES-KOH pH 8.0, 100 mM KCl, 2 mM EDTA, 0.1% NP40, 10% glycerol, 1x Protease inhibitor cocktail (25x, 11697498001, Sigma), 50 mM NaF, 1 mM PMSF, 1 mM DTT, 10 μg/ml TPCK) in a 1:4 pellet:buffer ratio. Samples were snap frozen in liquid nitrogen and thawed at 37 °C while agitating. Samples were sonicated for 30 sec and treated with 125 U Benzonase for 1 hour at 4 °C. After centrifugation at 16600 g for 30 minutes at 4 °C the supernatant was transferred to a fresh tube and the protein concentration was measured by Bradford.

### Affinity purification of protein complexes

Lysates were used at 2 mg/ml for immunoprecipitation (2 mg for western blot, 12 mg for mass spectrometry). 100 μl lysate was used as input sample. The remaining sample was incubated with Strep-Tactin XT magnetic beads (IBA) or Dynabeads™ (Thermo Fisher Scientific, 14311D) coupled with a NUP98 antibody (Abcam, ab50610) for 1 hour at 4 °C while rotating. Bead-protein complexes were washed three times with AP-MS buffer and eluted in 2% SDS buffer (50 mM HEPES pH8.0, 150 mM NaCl, 5 mM EDTA, 2% SDS). Eluates were subsequently used for western blot analysis or submitted to tryptic digestion and LC-MS/MS analysis.

### Biotinylated isoxazole-mediated precipitation

The assay was performed as previously described ^36^ with slight modifications. 10-20×10^6^ cells were resuspended in 1 ml ice-cold lysis buffer (50 mM HEPES-NaOH pH7.4, 150 mM NaCl, 0,1% NP40, 1 Mm EDTA, 2,5 mM EGTA, 10% Glycerol, 1x Protease inhibitor cocktail (25x), 50 mM NaF, 1 mM PMSF, 1 mM DTT, 10 μg/ml TPCK) and incubated at 4 °C for 30 minutes while rotating. After centrifugation at 16600 g for 30 minutes at 4 °C the supernatant was transferred to a fresh tube. After removing 50 μl of whole cell extract (input sample), the remaining sample was split in three fractions and treated with different concentrations of biotinylated isoxazole (10 mM stock in DMSO). Samples were incubated for 1 hour at 4 °C while rotating. Precipitated proteins were pelleted at 16600 g and washed three times with lysis buffer. Washed precipitates were resuspended in Laemmli buffer (western blot) or directly submitted to sample preparation for LC-MS/MS.

### Western blotting

Western blotting was performed according to standard laboratory protocols. Antibodies used were: anti-HA.11 (BioLegend, 901513; 1:2000, RRID: AB_2565335), anti-alpha Tubulin (Abcam, ab7291; 1:5000, RRID: AB_2241126), anti-beta Actin (Cell Signaling, 4967S; 1:5000, RRID: AB_330288), anti-HSC70 (Santa Cruz, sc-7298; 1:10000, RRID: AB_627761), anti-NUP98 (Cell Signaling, #2288; 1:1000, RRID: AB_561204), anti-RAE1 (Cell Signaling, sc-374261; 1:1000, RRID: AB_11008069).

Secondary antibodies used were: sheep anti-mouse HRP (GE Healthcare Austria GmbH & Co OG, NA931V; 1:10000, RRID: AB_772210), goat anti-rabbit HRP (Cell Signaling, 7074; 1:10000, RRID: AB_2099233).

### Filter Aided Sample Preparation and stage tip purification for AP-MS

100 μl of eluted protein complexes were used for Filter Aided Sample Preparation (FASP) as reported by Wisniewski et al.^61^. Samples were reduced by adding Dithiothreitol (DTT) to a final concentration of ~83.3 mM, incubated for 5 min at 99 °C before cooling down to room temperature. 400 μl 8 M urea in 100 mM Tris-HCl pH 8.5 (UA buffer) was added to the sample, mixed and sample transferred to a UA buffer-washed Microcon-30 kDa Centrifugal Filter Unit (Millipore, MRCF0R030). After an additional wash step with UA buffer, 100 μl of iodoacetamide (50 mM) was added to the filter unit, mixed at 800 rpm for 1 min before incubating for 30 min in the dark at room temperature. Filters were then washed three times with UA buffer, followed by three washes with 50 mM Triethylammonium bicarbonate (TEAB pH 8.5) buffer. To digest proteins, 1 μg trypsin (Promega, V511X) in 50 mM TEAB buffer was added and samples were incubated overnight at 37 °C. After additional washing steps with 50 mM TEAB buffer, digested peptides were eluted with 50 μl of 0.5 M NaCl. Samples were acidified with 5 μl 30% trifluoroacetic acid (TFA) and subsequently loaded onto in-house fabricated C18 stage tips. Desalted peptides were eluted using 0.4% formic acid with 90% acetonitrile. After vacuum centrifugation, peptides were reconstituted in 5% formic acid and submitted to LC-MS/MS analysis.

### In-solution digestion for biCon-MS

B-isox precipitates were resuspended in 8M urea in 100 mM TEAB buffer, pH 8 and proteins reduced with a final concentration of 50 mM DTT and incubated at 60°C for 1 hour. After cooling down to room temperature, reduced cysteins were alkylated with iodoacetamide at a final concentration of 55 mM for 30 min in the dark. Prior to tryptic digestion, urea concentration was diluted with 100 mM TEAB buffer pH 8 to 1.5 M and samples were digested with 1.25 μg of trypsin overnight at 37°C. Peptides were cleaned up by acidifying the samples to a final concentration of 1% TFA prior to performing solid phase extraction using C18 SPE columns (SUM SS18V, NEST group, USA) according to the manufacturer’s instructions. Peptides were eluted using two times 50 μl 90% Acetonitrile, 0.4% formic acid, organic solvent removed in a vacuum concentrator before reconstituting in 26 μl of 5% formic acid (Suprapur, MERCK KgaA, Germany).

Liquid chromatography mass spectrometry was performed on a hybrid linear trap quadrupole (LTQ) Orbitrap Velos mass spectrometer (ThermoFisher Scientific, Waltham, MA) or a Q Exactive™ Hybrid Quadrupole-Orbitrap (ThermoFisher Scientific, Waltham, MA) coupled to an Agilent 1200 HPLC nanoflow system (Agilent Biotechnologies, Palo Alto, CA) via nanoelectrospray ion source using a liquid junction (Proxeon, Odense, Denmark). Tryptic peptides were loaded onto a trap column (Zorbax 300SB-C18 5 μm, 5 × 0.3 mm, Agilent Biotechnologies) at a flow rate of 45 μL/min using 0.1% TFA as loading buffer. After loading, the trap column was switched in-line with a 75 μm inner diameter, 25 cm analytical column (packed in-house with ReproSil-Pur 120 C18-AQ, 3 μm, Dr. Maisch, Ammerbuch-Entringen, Germany). Mobile-phase A consisted of 0.4% formic acid in water and mobile-phase B of 0.4% formic acid in a mix of 90% acetonitrile and 9.610% water. The flow rate was set to 230 nL/min and a 90 min gradient used (4% to 30% solvent B within 81 min, 30% to 65% solvent B within 8 min and, 65% to 100% solvent B within 1 min, 100% solvent B for 6 min before equilibrating at 4% solvent B for 18 min). For the MS/MS experiment, the LTQ Orbitrap Velos mass spectrometer was operated in data-dependent acquisition (DDA) mode with the 15 most intense precursor ions selected for collision-induced dissociation (CID) in the linear ion trap (LTQ). MS^1^-spectra were acquired in the Orbitrap mass analyzer using a scan range of 350 to 1,800 m/z at a resolution of 60,000 (at 400 m/z). Automatic gain control (AGC) was set to a target of 1 × 10^6^ and a maximum injection time of 500 ms. MS^2^-scans were acquired in parallel in the linear ion trap with AGC target settings of 5 × 10^4^ and a maximum injection time of 50 ms. Precursor isolation width was set to 2 Da and the CID normalized collision energy to 30%. The Q Exactive™ MS was operated in a Top10 DDA mode with a MS^1^ scan range of 375 to 1,650 m/z at a resolution of 70,000 (at 200 m/z). Automatic gain control (AGC) was set to a target of 1 × 10^6^ and a maximum injection time of 55 ms. MS^2^-spectra were acquired at a resolution of 15,000 (at 200 m/z) with AGC settings of 1 × 10^5^ and a maximum injection time of 110 ms. Precursor isolation width was set to 1.6 Da and the HCD normalized collision energy to 28%. The threshold for selecting MS^2^ precursor ions was set to ~2,000 counts for both instruments. Dynamic exclusion for selected ions was 30 s. A single lock mass at m/z 445.120024 was employed (Olsen, de Godoy et al., 2005). All samples were analyzed in technical duplicates. Xcalibur version 2.1.0 SP1/Tune2.6.0 SP3 and XCalibur version 4.1.31.9 Tune 2.9.2926 were used to operate the LTQ Orbitrap Velos or Q Exactive MS instrument, respectively.

### MS data analysis with SearchGUI / PeptideShaker (Figure 1 and Figure 2)

We used msconvert^62^ from the ProteoWizard^62^ toolkit to convert raw data files to mgf format. These mgf files were then processed with SearchGUI (version 3.2.20)^63^ with the xtandem, myrimatch, ms_amanda, msgf, omssa, comet and tide search engines, against the human Swiss-Prot^64^ database (01.2017) extended with the Strep-HA-tag, Strep-Tactin and trypsin protein sequences. As post-translational modifications we configured fixed carbomidomethylation of C and variable oxidation of M. The decoy database was generated within SearchGUI. Results were then analyzed with PeptideShaker (version 1.16.15) ^65^.

### MS data analysis with Proteome Discoverer (Figure 4 and Figure 5)

Acquired raw data files were processed using the Proteome Discoverer 2.2.0.388 platform, utilising the database search engine Sequest HT. Percolator V3.0 was used to remove false positives with a false discovery rate (FDR) of 1% on peptide and protein level under strict conditions. RAW data was recalibrated prior to Sequest HT searches using full tryptic digestion against the human SwissProt database v2017.06 (20,456 sequences and appended known contaminants) with up to one miscleavage site. Oxidation (+15.9949 Da) of methionine, acetylation (+42.010565 Da) of lysine and protein N-terminus and phosphorylation (+79.966331 Da) of serine, threonine and tyrosine were set as variable modifications, whilst carbamidomethylation (+57.0214 Da) of cysteine residues was set as fixed modifications. Data was searched with mass tolerances of ±10 ppm and 0.025 Da on the precursor and fragment ions, respectively. The ptmRS node was used for additional validation of posttranslational modifications. Results were filtered to include peptide spectrum matches (PSMs) with Sequest HT cross-correlation factor (Xcorr) scores of ?1, ptmRS scores of ?75 and protein groups including ?2 peptides. For calculation of protein amounts, the Minora Feature Detector node and Precursor Ions Quantifier node, both integrated in Thermo Proteome Discoverer were used. Automated chromatographic alignment and feature linking mapping were enabled. Precursor abundance was calculated using intensity of peptide features including only unique peptide groups and excluding phosphorylated and acetylated peptides. To equalize total abundance between different runs, protein abundance values were normalized using the total peptide amount approach. No computational missing value imputation was applied to fill gaps. For statistical analysis a nested (paired) approach was applied using pairwise ratio calculation and background-based ANOVA statistical testing. Pairwise ratio calculation was chosen to make the analysis less sensitive towards missing values. Background-based ANOVA uses the background population of ratios for all peptides and proteins in order to determine whether any given single peptide or protein is significantly changing relative to that background (as stated in the manual of Proteome Discoverer 2.2, Thermo Fisher Scientific, Waltham, MA). Adjusted p-values are calculated using the Benjamini-Hochberg method.

### Protein network analysis

Protein networks were illustrated with Cytoscape 3.6.1^66^. Annotated protein-protein interactions were obtained from the STRING database using the StringApp 1.4.2 in Cytoscape with a confidence score cutoff of 0.4. Reactome clustering for protein networks was performed using the Reactome FI 7.1.0 application and executed with default conditions. Gene Ontology networks were generated with ClueGO v2.5.4^67^ application with a P-value cutoff of 0.05.

### O/E ratios of binned PScores (Figure 3)

PScores of individual proteins found within each Gene Ontology term were grouped into 6 bins ranging from <0 to >4 (*observed* distribution). The same was done for the PScores of the human proteome^35^ (*expected* distribution). Next, the ratio between observed and expected frequencies were calculated for each bin and illustrated as a heatmap. The heatmap was generated with the heatmap.2 function from the R-package gplots (version 3.0.1)^68^.

### Group size dependent testing statistic (GSDTS)

GSDTS tests a list of values x against a larger universe y of values (x < y) by sub-sampling z-times a list of length of x out of the universe. Each sub-sampled list is checked against the already created lists to ensure unique lists to satisfy “sub-sampling without replacement”. The lists x and y as well as the number of sub-sampling repeats z need to be provided by the user. As result the sum, mean or median of all sub-sampling results are plotted as histogram and the respective sum, mean or median of list x is then indicated by a red bar in the histogram. The script is implemented in R and available at: https://github.com/Edert/R-scripts For the histograms shown, we applied GSDTS with a z of 10,000 to obtain 10,000 subsampled lists of length x of the universe y. As universe y we chose the whole human proteome ^35^ and x was chosen to match the length of the query list. We then used the mean to plot the histogram and for indicating the tested list as a red line.

### RNA-seq analysis for fetal liver cells

Cells were retrovirally transduced and FACS sorted for mCherry+ cells. After expansion for seven days, RNA was isolated according to the standard protocol using the RNeasy Mini Kit (QIAGEN). Sequencing libraries were generated using the QUANT™seq 3’ mRNA-Seq Library Prep Kit (Lexogen). Pooled libraries were processed at the Vienna Biocenter Next Generation Sequencing Facility (Vienna, Austria) and sequenced using single read 70bp chemistry on a NextSeq550 instrument (Illumina).

### RNA-seq data analysis

The quality of the raw sequence files was checked with FastQC (version 0.11.4)^69^. Based on this the quality trimming and filtering and length filtering was applied using PRINSEQ-lite (version 0.20.4)^70^. Remaining reads were aligned against the mouse reference genome (GRCm38/mm10) by STAR (version 2.5.0b)^71^. Final processing was carried out with SAMtools (version 1.7)^72^ and counts per gene were obtained by featureCounts (version 1.6.0)^73^. Differential gene expression analysis and normalization prior to visualization and clustering was then performed with DESeq2 (version1.22.2)^74^. Batch-correction for two library preparation and sequencing runs was done with ComBat^75^ from the sva package^76^. For the PCA the top 500 genes with the highest variance in their normalized expression were used.

The heatmap of shared regulated genes was generated with the heatmap.2 function from the R-package gplots (version 3.0.1)^68^ using Pearson correlation as distance measure and ward.D clustering.

## Data availability

LC-MS/MS data will be deposited into PRIDE under the accession number XXX RNA-seq data will be deposited into GEO under the accession number GEO: YYY

## Lead Contact and Materials Availability

All unique/stable reagents generated in this study are available from the Lead Contact with a completed Materials Transfer Agreement.

## Acknowledgments

We thank all members of the Grebien laboratory for stimulating discussions and A. Orlova for help with QUANT-seq experiments. A. Spittler, G. Hofbauer (Core Facility Flow Cytometry, Medical University of Vienna), S. Fajmann and P. Jodl (Institute of Pharmacology and Toxicology, University of Veterinary Medicine Vienna) are acknowledged for cell sorting. This research was supported using resources of the VetCore Facility (Imaging) of the University of Veterinary Medicine Vienna. Next Generation Sequencing was performed at the VBCF NGS Unit (www.viennabiocenter.org/facilities). In addition, we thank all members of the Proteomics and Metabolomics Facility (CeMM Research Center for Molecular Medicine of the Austrian Academy of Sciences) for the proteomics analysis. J.S. is a recipient of a DOC Fellowship of the Austrian Academy of Sciences at the University of Veterinary Medicine Vienna. This work was supported by a grant from the European Research Council under the European Union’s Horizon 2020 research and innovation programme (grant agreement n° 636855/StG) to F.G.

## Author Contributions

Conceptualization S.T-Z, F.G.L and F.G.; Methodology S.T-Z, J.S., F.G.L and F.G.; Formal Analysis S.T-Z., T.E. and A.C.M.; Investigation S.T-Z., J.S., T.H., N.K., K.P. and E.H.; Data Curation T.E. and A.M.; Writing - Original Draft S.T-Z. and F.G.; Visualization S.T-Z., T.E., T.H., N.K. and F.G.; Supervision F.G.; Project Administration S. T-Z. and F.G.; Funding Acquisition F.G.

## Supplemental Information titles and legends

**Figure S1.**
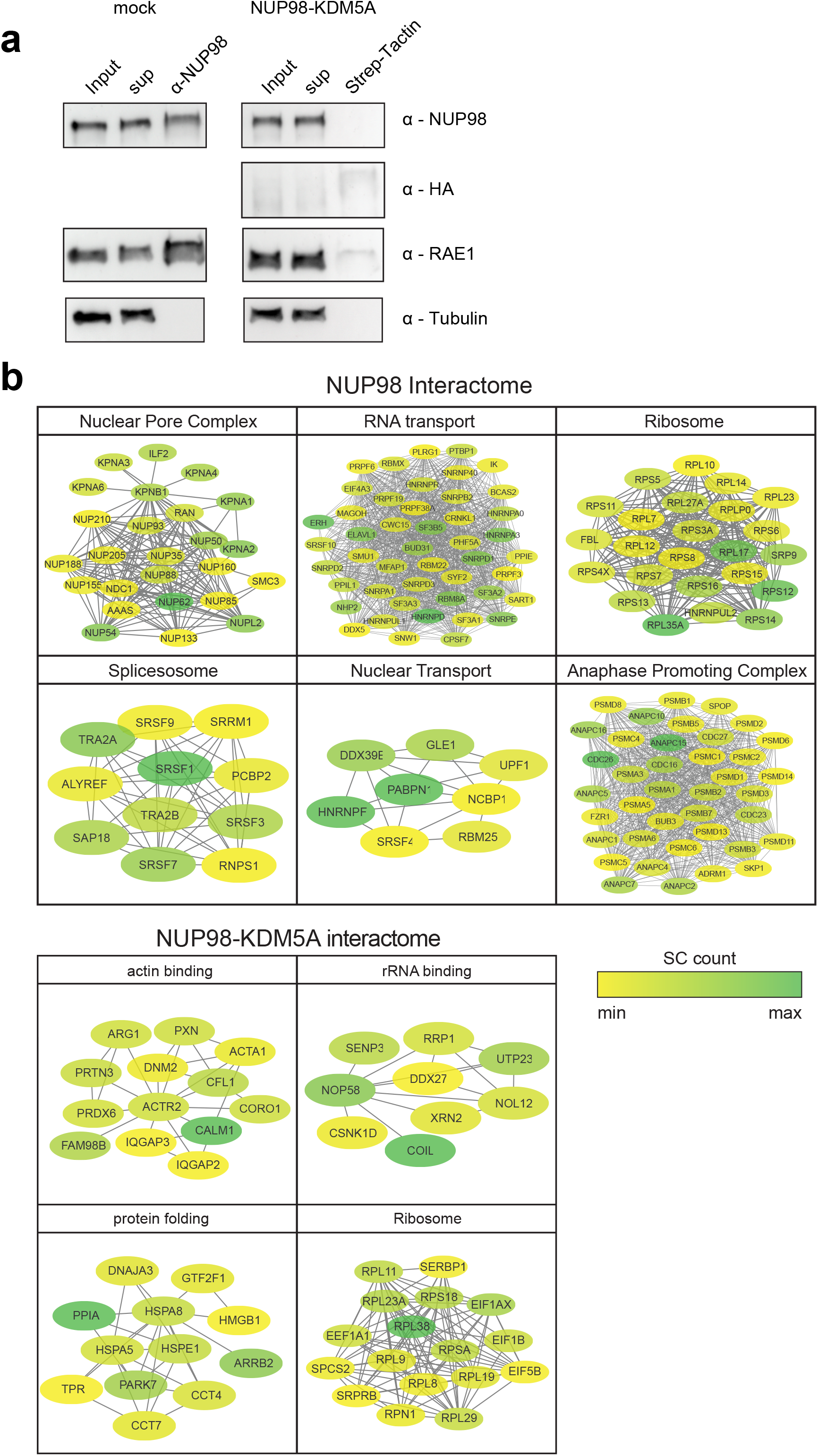
Immunoprecipitation of endogenous NUP98 and affinity purification of NUP98-KDM5A coupled to LC-MS/MS, Related to Figure 1. (a) Western blot analysis of mock and NUP98-KDM5A-expressing HL-60 cells. Mock lysates were incubated with NUP98 antibody conjugated to magnetic beads and NUP98-KDM5A lysates were incubated with magnetic Strep-Tactin beads. Input, supernatant and pull down were detected with anti-NUP98, anti-HA, anti-RAE1 and anti-Tubulin. (b) STRING networks of individual subcomplexes identified by Gene Ontology for Biological Processes. Color gradient indicates abundance in mass spectrometry.

**Figure S2.**
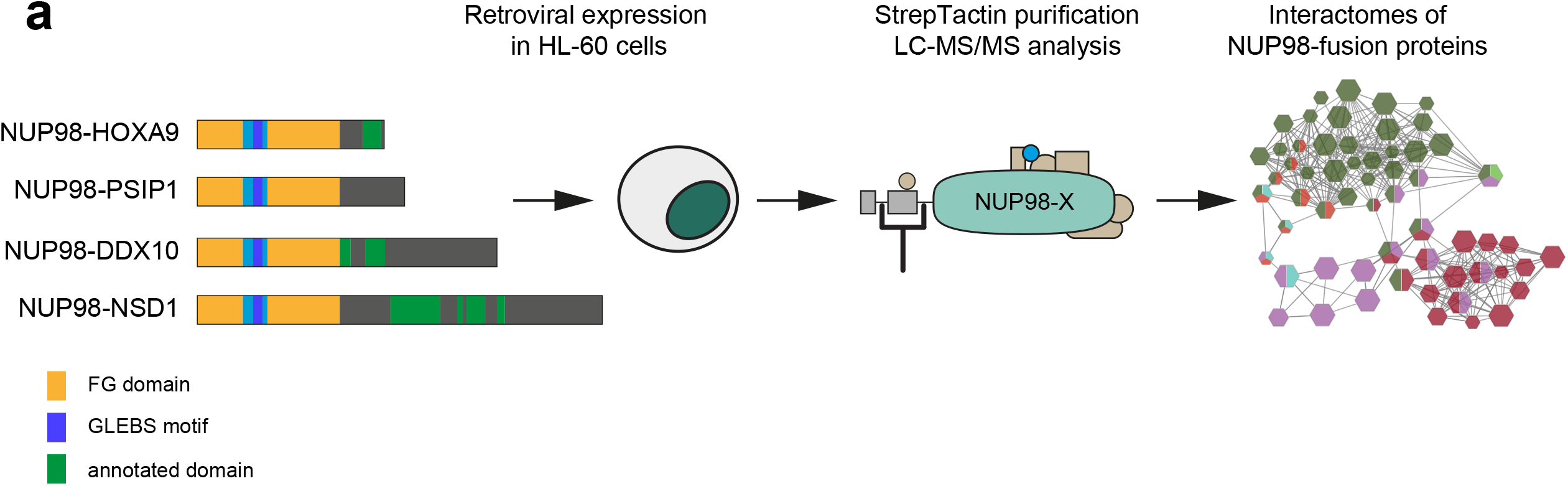
AP-MS of a diverse set of NUP98-fusion proteins, Related to Figure 2. (a) Schematic representation of NUP98-fusion proteins and stable transduction of HL60 cells. Protein complex pulldowns were performed using Strep-Tactin followed by LC-MS/MS analysis.

**Figure S3.**
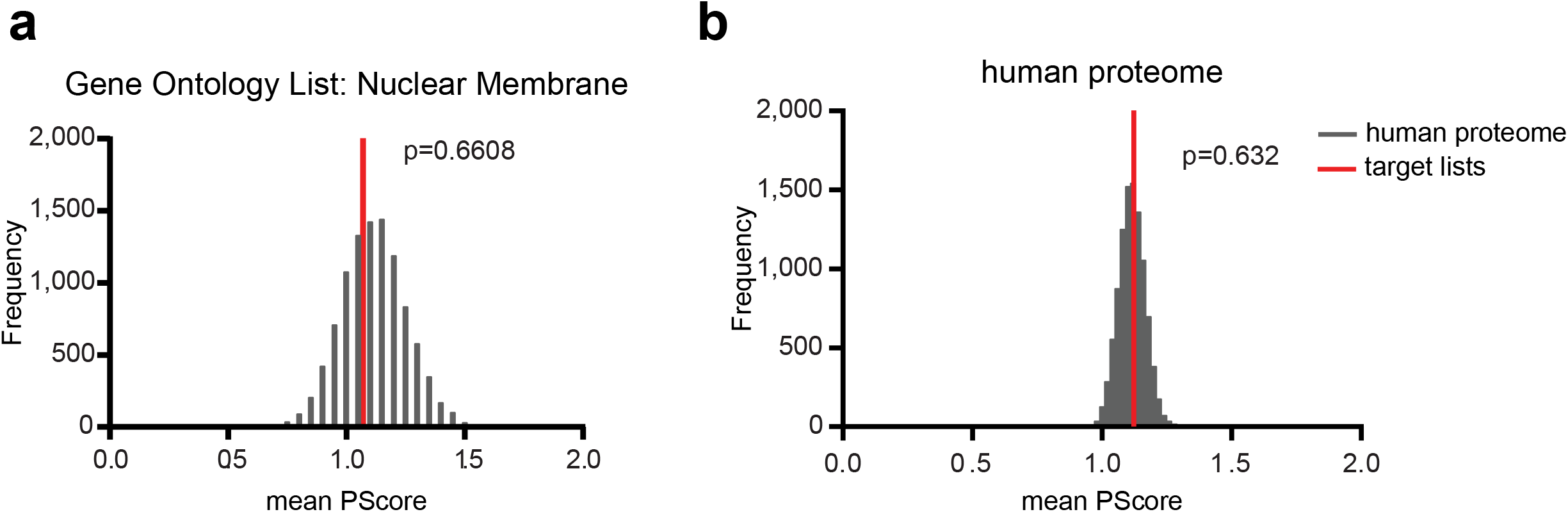
PScore analysis of different protein lists, Related to Figure 3. (a) Mean PScore of the Gene Ontology list “Nuclear Membrane” was compared to a set of randomly subsampled lists of the human proteome with the same size. (b) Mean PScore of the human proteome was compared to a set of randomly subsampled lists of the human proteome with the same size. P-values were calculated using the Kolmogorov-Smirnov test.

**Figure S4.**
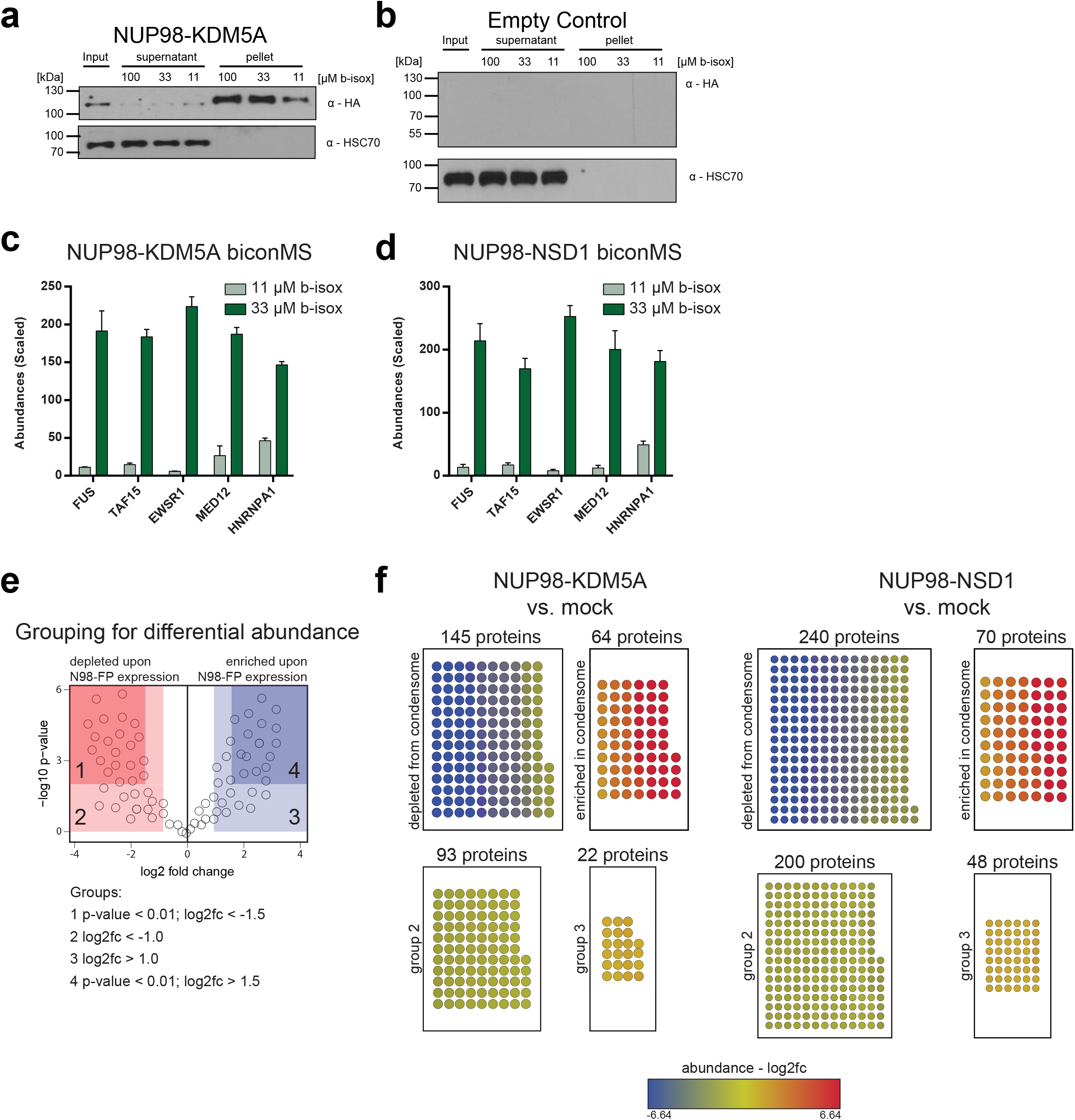
biCon-MS for NUP98-KDM5A and NUP98-NSD1, Related to Figure 5. (a) Western blot analysis of NUP98-KDM5A-expressing NIH-3T3 cell lysates treated with 11 μM, 33 μM or 100 μM b-isox. Dose-dependent precipitation was investigated for NUP98-KDM5A and HSC70. (b) Western blot analysis of NIH-3T3 cell lysates treated with 11 μM, 33 μM or 100 μM b-isox. Dose-dependent precipitation was investigated for HSC70. (c) Scaled abundance for proteins previously implicated in the formation of biomolecular condensates identified by biCon-MS from lysates of HL-60 cells expressing NUP98-KDM5A. (d) Scaled abundance for proteins previously implicated in the formation of biomolecular condensates identified by biCon-MS within lysates of HL-60 cells expressing NUP98-NSD1. (e) Schematic illustration of enriched/depleted proteins identified in fusion protein biCon-MS compared to mock HL-60 cells (f) Enrichment of proteins that exhibit dose-dependent precipitation upon expression of NUP98-KDM5A and NUP98-NSD1 as compared to mock HL-60 cells based on abundance in biCon-MS analysis. Enriched/depleted proteins in NUP98-fusion protein condensates are illustrated as nodes and are colored according to calculated fold changes. Depleted cutoff: log2(fc) < -1.5 and P-value < 0.01 Enriched cutoff: log2(fc) > 1.5 and P-value < 0.01

**Figure S5.**
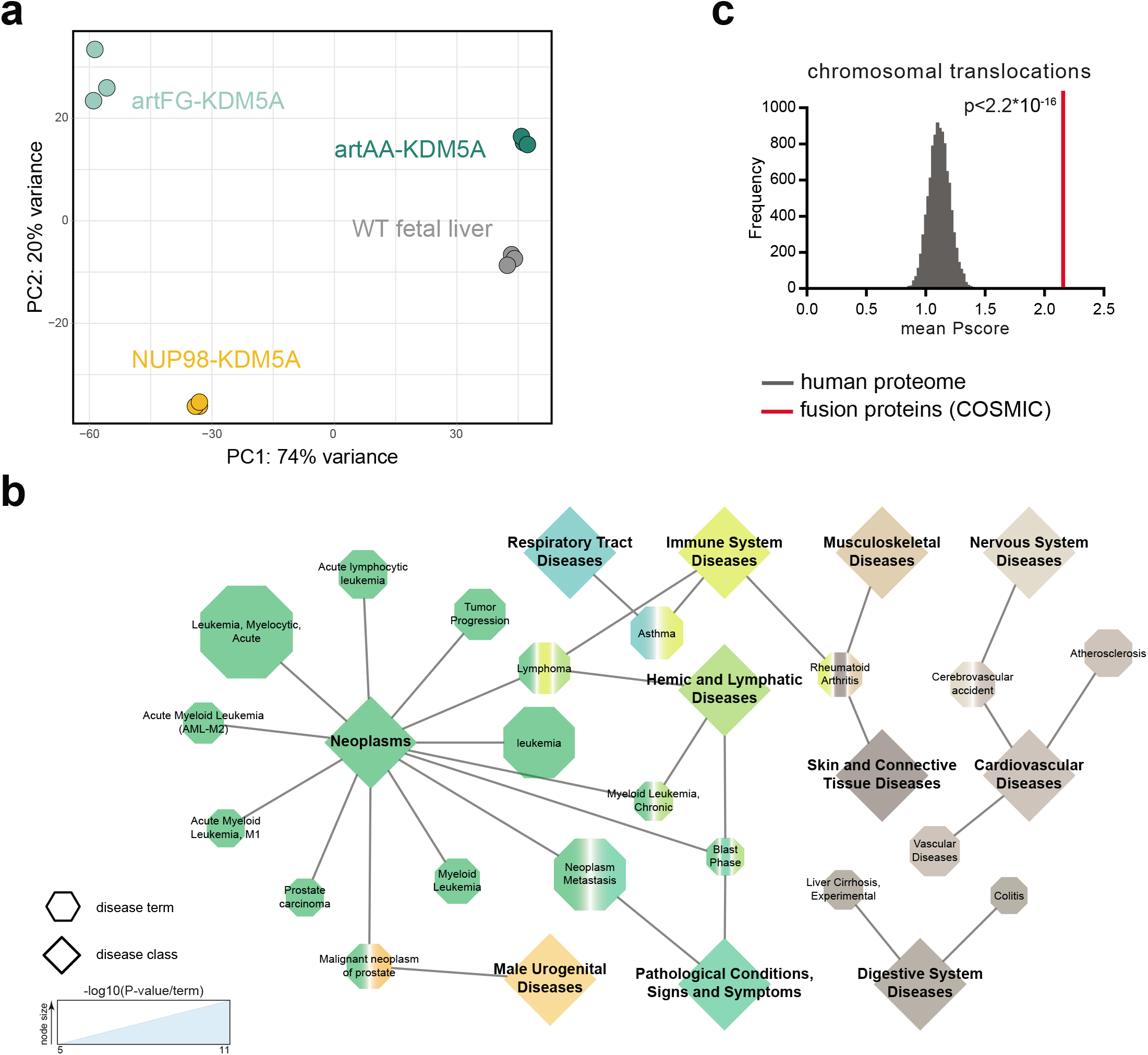
Related to Figure 6. (a) Principal component analysis of individual RNA-seq replicates. (b) 761 differentially regulated genes of NUP98-KDM5A and artFG-KDM5A were group using DisGeNET according to related diseases. Most significant disease terms are illustrated as hexagon sized according to their P-value. Corresponding disease classes are shown as diamond connected by edges to diseases, defined by DiGeNET. (c) Mean PScore for all proteins involved in cancer gene fusions listed in COSMIC was compared to a list of randomly subsampled lists of the human proteome with the same size. P-value was calculated using the Kolmogorov-Smirnov test.

**Supplementary Table 1. Mass spectrometry of endogenous NUP98 and NUP98-KDM5A**

**Supplementary Table 2. Mass spectrometry of NUP98-HOXA9, NUP98-PSIP1, NUP98-DDX10, NUP98-NSD1 and spectrum counts for core interactors**

**Supplementary Table 3. Mass spectrometry of biCon-MS for HL60 cells (mock, NUP98-KDM5A and NUP98-NSD1)**

